# Validation of EEG data assimilation-based prefrontal excitation-inhibition balance estimation using TMS–EEG

**DOI:** 10.1101/2025.03.22.644706

**Authors:** Hiroshi Yokoyama, Yoshihiro Noda, Masataka Wada, Mayuko Takano, Keiichi Kitajo

## Abstract

The excitation and inhibition (E/I) balance of neural circuits is a crucial index reflecting neurophysiological homeostasis. Although several cutting-edge methods have been established to assess E/I balance in an intact brain, they have inherent limitations, such as difficulties in tracking changes in E/I balance over time. To tackle this issue, we have proposed neural-mass-model-based tracking of the brain states using a data assimilation (DA) scheme in our previous work. However, although we verified that sleep-dependent changes in E/I balance can be estimated from electroencephalography data, the neurophysiological validity of the method was not evaluated. Therefore, in the current study, we directly compared estimated E/I states based on the DA methods with the concurrent transcranial magnetic stimulation and electroencephalography (TMS-EEG) based methods. The results showed that the E/I changes estimated by the DA-based method correlated significantly with E/I in the dorsolateral prefrontal cortex, as indexed by TMS-evoked EEG. These findings indicate that our proposed method can estimate neurophysiologically valid changes in E/I balance.

## Introduction

Maintaining an appropriate excitation/inhibition (E/I) balance in neural circuits is essential for normal brain function. The breakdown of the system of E/I balance can lead to various neuropsychiatric disorders. For example, E/I imbalance caused by GABAergic inhibitory and/or glutamatergic excitatory dysfunctions constitutes the pathophysiological neural basis of various neuropsychiatric disorders, including schizophrenia, bipolar disorder, major depressive disorder, autism spectrum disorder, and Alzheimer’s disease [1–9]. Moreover, several postmortem brain studies have reported abnormal expression level of GABA-related genes and molecules in the prefrontal cortex of patients with neuropsychiatric disorders (mainly major depressive disorder, autism spectrum disorder, and attention-deficit/hyperactivity disorder) [10, 11]. Therefore, insight into normal and abnormal brain function in relation to neurophysiological E/I balance is critical for elucidating the pathophysiology of neuropsychiatric disorders.

To date, two existing measurement methods are typically used to evaluate E/I balance in the human brain: magnetic resonance spectroscopy (MRS) and simultaneous recording of electroencephalography with transcranial magnetic stimulation (TMS–EEG). However, these methods have inherent limitations, including difficulties in tracking changes in E/I balance over time and in assessing E/I balance over short time intervals. To address these issues, in our previous work [12], we proposed and implemented a method for tracking temporal changes in E/I balance based on the neural-mass (NM) model from human scalp electroencephalography (EEG) using the data assimilation (DA) method. In this previous study, the NM model, which contains parameters regarding E/I synaptic gains, was directly assimilated to the observed EEG data using the Ensemble Kalman Filter (EnKF) based method [12]. As a result, this proposed method can reconstruct the temporal changes in E/I balance from human scalp EEG based on the estimated E/I parameters of the NM model in a data-driven manner. The validity of the proposed method was confirmed by showing that it could assess sleep-dependent changes in E/I balance from human EEG data on the sub-second scale. The tendency of such sleep-dependent changes was consistent with the experimental hypothesis reported on sleep-dependent changes in E/I balance in a previous human MRS study [13]

However, MRS is limited to measuring the total concentration of neurotransmitters within a specific brain region and cannot accurately assess GABA and glutamine concentrations in extrasynaptic areas [14]. Moreover, some studies highlighted that total GABA concentrations observed with MRS are less sensitive to GABAergic inhibitory activity because of the above-mentioned smaller accuracy for estimating total neurotransmitter concentrations [14–16]. Given these methodological limitations of MRS, our recently proposed DA method [12] indicates that it is possible to evaluate changes in sleep-dependent E/I balance over time, consistent with previous findings [13]. However, the neurophysiological validity of this method remains unconfirmed.

To address this remaining issue, we aimed to validate our proposed method by directly comparing the DA-based estimation of E/I balance with TMS neurophysiological indices using TMS-EEG, which can physiologically evaluate GABAergic inhibitory and glutamatergic excitatory functions [3, 5, 17, 18]. In this study, we used the following two types of EEG data collected from the same participants: (1) short-interval intracortical inhibition (SICI, an index reflecting GABA_A_ receptor-mediated neurophysiological function) and intracortical facilitation (ICF, an index reflecting glutamate N-methyl-D-aspartate receptor-mediated neurophysiological function), both derived from the TMS-EEG method, and (2) resting-state EEG. Each dataset was used to assess TMS-EEG-based and EEG-DA-based estimates of E/I balance. Both estimation results were directly compared to validate the electrophysiological accuracy of our DA-based method. In addition, to improve the computational cost of E/I estimation [12], we modified our proposed EEG-DA method by combining our previously proposed algorithms: variational Bayesian noise-adaptive constrained EnKF (vbcEnKF [12, 19–27]) and Ensemble Transform Kalman Filter (ETKF) [28].

## Results

### Study overview

Recently, we proposed a sequential DA-based method to estimate E/I changes from scalp EEG; however, sufficient neurophysiological accuracy was not confirmed [12]. To address this issue, we have directly compared intracortical E/I evaluation results between DA-based and TMS-EEG-based methods (see Fig. 1).

**Fig. 1.**
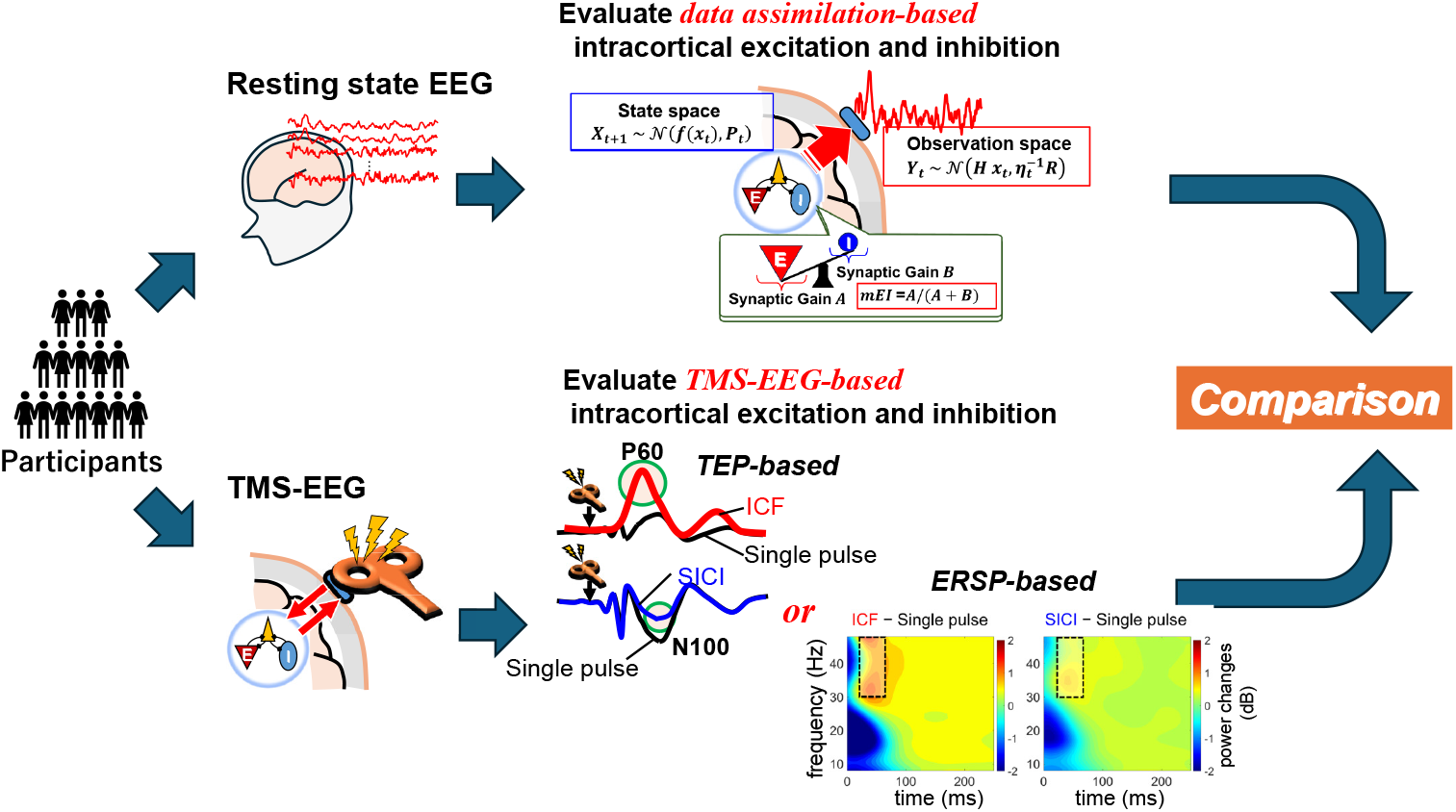
Study overview: electrophysiological validation of excitation/inhibition (E/I) estimation based on electroencephalography and data assimilation (EEG-DA) by comparison with transcranial magnetic stimulation and electroencephalography (TMS-EEG). Participants took part in two experiments: resting-state EEG and concurrent TMS-EEG recordings. For resting-state EEG recordings, EEG signals were recorded for 5 min from the scalp of each participant. Our proposed DA-based E/I estimation method was applied to the EEGs obtained from five electrodes around left dorsolateral prefrontal cortex (L-DLPFC). In the DA-based analysis, the five EEG time series were individually fitted with a neural-mass model, and resulting time series of the E and I gain parameters, *A* and *B*, were temporally averaged for each electrode. Averaged *A* and *B* values were directly compared with the TMS-EEG-based E/I estimation. In the concurrent TMS-EEG recordings, the EEGs were recorded from the scalp of each participant while applying TMS to L-DLPFC. The TMS-EEG-based E/I estimation was assessed based on two approaches using either conventional evoked potential (e.g., N100 and P60) or event-related spectral perturbation (ERSP). Each approach was accessed by intracortical facilitation (ICF) and short-interval intracortical inhibition (SICI) using TMS-evoked potential (TEP) and ERSP-based evaluation indices, respectively. ICF and SICI indicate a constant dynamic range of glutamatergic-mediated excitation and GABAergic-mediated inhibition in the brain.

### TMS-EEG based E/I estimation

In this study, we used either TMS-evoked potential (TEP) [18, 29–31] or event-related spectral perturbation (ERSP) [32, 33] as TMS-EEG-based E/I evaluation indices (Fig. 1). In the TEP-based method, the TEP component regarding GABAergic-mediated SICI was calculated by subtracting the negative peak of TMS-triggered potentials between single and paired pulse around 70–130 ms intervals after TMS onset (called N100), i.e., *SICI* = *N*100_*single*_ −*N*100_*paired*_. Additionally, the TEP component regarding glutamatergic-mediated ICF was calculated by subtracting the positive peak of TMS-triggered potentials between single and paired pulse around 30–80 ms intervals after TMS onset (called P60), i.e., *ICF* = *P*60_*paired*_ − *P*60_*single*_. To obtain SICI and ICF, inter-stimulus intervals in the paired-pulse TMS of 1–5 ms and 10–15 ms were selected, respectively. For the ERSP-based method, the ERSP of SICI was calculated by subtracting the gamma power obtained from the SICI from that obtained from single-pulse TMS across five electrodes around the left dorsolateral prefrontal cortex (L-DLPFC).

Similarly, the ERSP of ICF was calculated by subtracting the gamma power obtained from single-pulse TMS from that obtained from the ICF paradigm.

Fig. 2 displays the relationships between the ICF and SICI components of each participant group. Correlation analysis was conducted using the opensource MATLAB robust correlation analysis toolbox [34]. TMS-evoked EEG responses in both participant groups were significantly correlated with each other (group 1: TEP-based estimation, Spearman’s rank correlation coefficient *r*_*sp*_ = 0.3990, *p* = 0.0160; group 2: ERSP-based estimation, Spearman’s rank correlation coefficient *r*_*sp*_ = 0.3388, *p* = 0.0335; Fig. 2). This indicates that E/I balance between glutamatergic and GABAergic functions maintains a consistent condition among participants. As these results (Fig. 2) were consistent with those of previous studies [30, 31], we can conclude that the E/I components estimated from the TMS-EEG experiments reliably reflect E/I features, independent of the evaluation method used (TEP or ERSP).

**Fig. 2.**
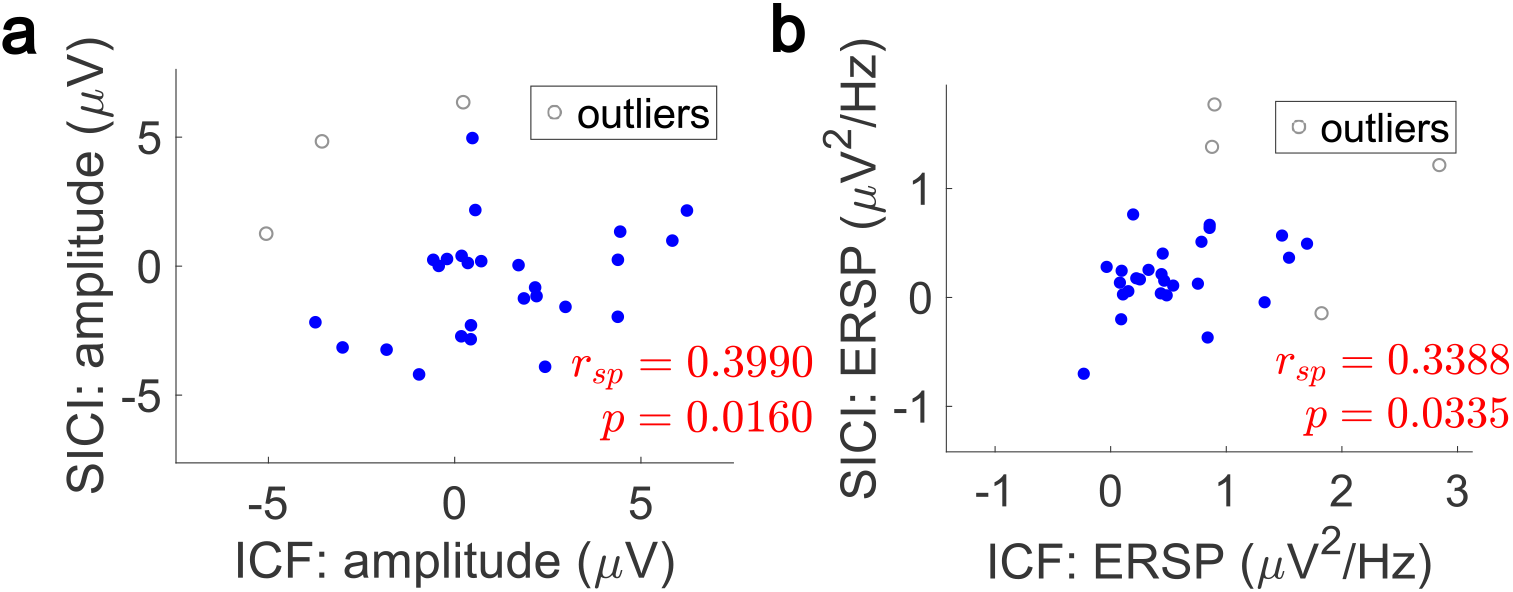
Estimations of excitation/inhibition based on transcranial magnetic stimulation-evoked potential (TEP) and event-related spectral perturbation (ERSP). **a**) The relationship between intracortical facilitation (ICF) and short-interval intracortical inhibition (SICI) components based on TEP amplitude. ICF was evaluated as the difference in P60 electroencephalography (EEG) amplitude between single- and paired-pulse responses around the left dorsolateral prefrontal cortex (DLPFC). The P60 component was calculated as the trial-averaged EEG signal around 60 ms after transcranial magnetic stimulation (TMS) onset. SICI was evaluated as the difference in N100 EEG amplitudes between single- and paired-pulse responses. The N100 peak was calculated as the trial-averaged EEG signal around 100 ms after TMS onset. **b**) The relationship between ICF and SICI components based on TMS-evoked changes in gamma frequency (30–48 Hz) power. The *r*_*sp*_ and *p* values indicate the skipped Spearman’s correlation coefficient and its *p*-value estimated by Robust Correlation Toolbox [34], respectively.

### DA-based E/I estimation

In the DA-based method, E/I features are evaluated based on the temporally averaged E and I gain parameters *A* and *B* in the NM model, estimated from resting-state EEG signals (Fig. 1). The time-varying changes in the parameters *A* and *B* were individually evaluated for each of five electrodes around L-DLPFC by directly assimilating the observed EEGs into the NM model using our proposed EEG-DA method. The resultant time-series values of *A* and *B* were temporally averaged for each of five electrodes. To estimate *A* and *B*, EEGs during the 5-minute resting-state task were directly assimilated into the NM model by using our proposed method. As mentioned in the Introduction, even though we recently proposed an EnKF-based DA method that enables tracking the time-varying changes in E/I balance indexed by the ratio of estimated E/I synaptic gain parameters, this method still has some issues, such as the computational cost of the EnKF algorithm [12, 35, 36] for the state estimation of the NM model. Therefore, to decrease the computational cost of model parameter estimation when assimilating the NM model into observed data, we modified this proposed DA-based E/I estimation method by combining the previously proposed method (vbcEnKF) with the ETKF algorithm. This algorithm can decrease the computational cost for state estimation in the NM model (see the Methods and [28] for more details). The numerical validation of this modified algorithm (that we called vbcETKF) is shown in the Supplementary Results.

By applying our modified DA method to the synthetic EEGs generated by the NM model with known parameters and E/I balance changes, we found that our method can correctly estimate the E/I gain parameters *A* and *B* and E/I balance changes with a number of ensemble member *N*_*ens*_ = 70. As EnKF is based on Sequential Monte Carlo sampling to estimate model parameters (see Methods and Supplementary Results for more details), the number *N*_*ens*_ is directly associated with the extent of the computational cost to estimate the NM model parameters in our DA method. However, by modifying our previous EnKF-based DA-method with ETKF, the method succeeded in drastically decreasing the required Nens compared with the previous EnKF-based method [12] (see Supplementary Results).

Before comparing the results between DA-based and TMS-EEG-based estimations of E/I balance, we assessed whether the estimated E/I gain parameters *A* and *B* in the NM model show significant correlations across participants in the same manner as in Fig. 2. The results of the E/I correlation analysis of the DA-based method are shown in Fig. 3. The DA-based E/I estimation was individually conducted in five EEG electrodes located around L-DLPFC, which is consistent with the electrodes to evaluate the ICF and SICI in the TMS-EEG-based method for E/I estimation. The E/I balance indexed by two estimated E/I gain parameters *A* and *B* using our DA method showed a significant correlation for all electrodes, independent of the participant groups (Fig. 3.).

**Fig. 3.**
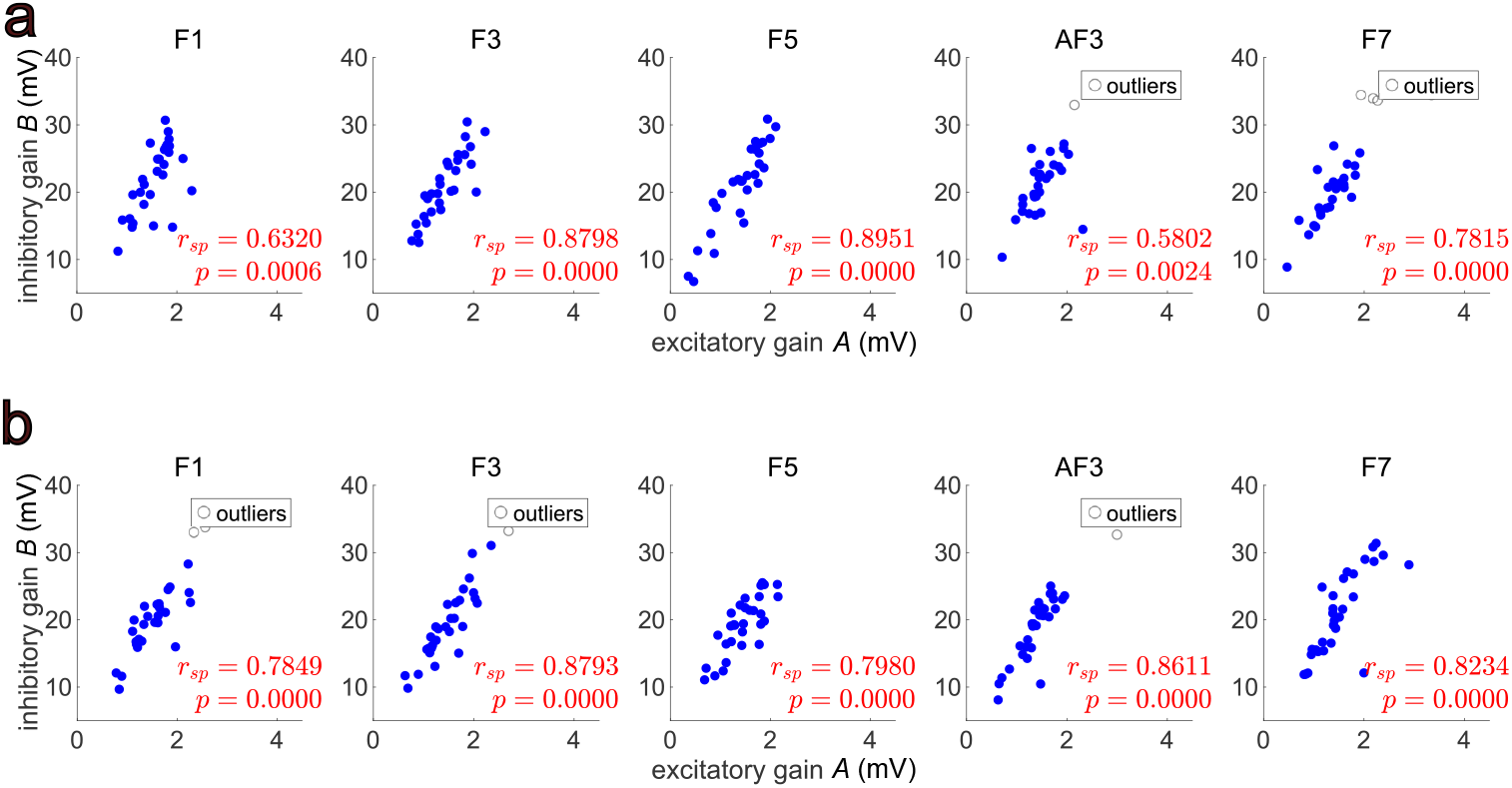
Relationships between data assimilation (DA)-based inhibitory and excitatory gains. **a, b**) **a, b**) Relationships between the excitation and inhibitory gain parameters *A* and *B* obtained from the DA of resting-state electroencephalography (EEG) data in group 1 (**a**) and group 2 (**b**). Each panel shows a scatter plot of correlation between the estimated excitation and inhibitory gain parameters (*A* and *B*) evaluated in EEGs obtained from five different electrodes (F1, F3, F5, AF3, and F7) around the left dorsolateral prefrontal cortex (DLPFC). The *r*_*sp*_ and *p*-values indicate the skipped Spearman’s correlation coefficient and its *p*-value evaluated by Robust Correlation Toolbox [34], respectively. The *p*-values were corrected by the Bonferroni method to avoid the multiple comparison effect.

Moreover, we found that the E/I balance estimated by the DA-based method shows an extremely strong correlation compared with the TMS-EEG-based method. In group 1 (Fig. 3a), the Spearman’s rank correlation coefficient was ≥ 0.58 in all electrodes (F1: *r*_*sp*_ = 0.6320, *p* = 0.0006, F3: *r*_*sp*_ = 0.8798, *p* = 0.0000, F5: *r*_*sp*_ = 0.8951, *p* = 0.0000, AF3: *r*_*sp*_ = 0.5802, *p* = 0.0024, F7: *r*_*sp*_ = 0.7851, *p* = 0.0000). Additionally, in group 2 (Fig. 3b), the correlation coefficient was ≥ 0.78 in all electrodes (F1: *r*_*sp*_ = 0.7849, *p* = 0.0000, F3: *r*_*sp*_ = 0.8793, *p* = 0.0000, F5: *r*_*sp*_ = 0.7980, *p* = 0.0000, AF3: *r*_*sp*_ = 0.8611, *p* = 0.0000, F7: *r*_*sp*_ = 0.8234, *p* = 0.0000).

These results indicate that the E/I balance indexed by the two estimated E/I parameters *A* and *B* using our DA method is sustained across participants as well as with the TMS-EEG-based method.

### Comparison of E/I estimations based on TMS-EEG and DA methods

We revealed that the E/I balance indexed by both TMS-EEG and DA-based methods was consistently sustained across participants (Figs. 2 and 3). This suggests that the DA-based method can detect electrophysiological valid features such as those that reflect the ICF and SICI. Therefore, to test the neurophysiological validity of our DA-based E/I estimation method, we directly compared the E/I estimation results of DA-based and TMS-EEG-based methods in groups 1 and 2 using the robust correlation toolbox (Figs. 4 and 5).

**Fig. 4.**
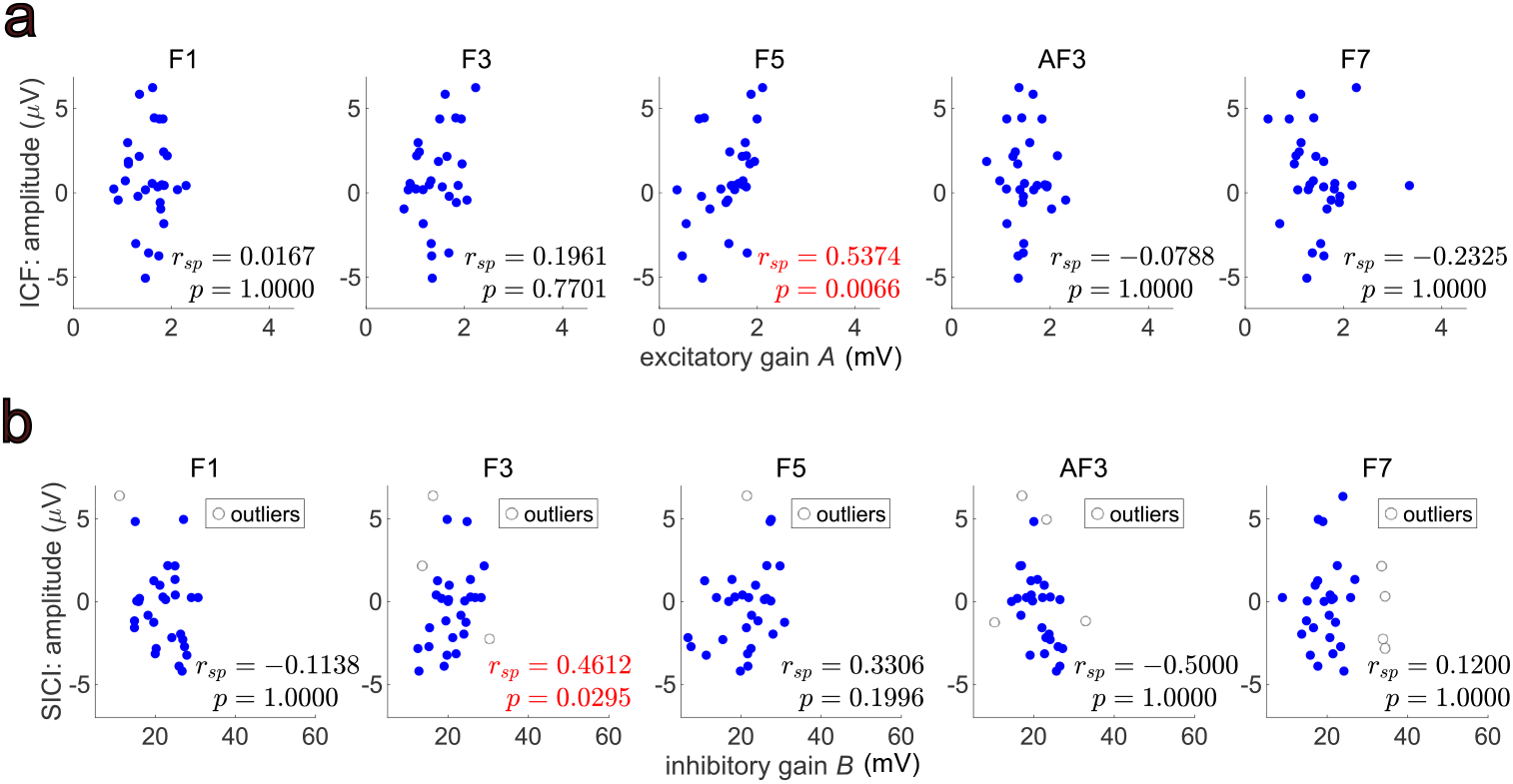
Excitation/inhibition estimations based on data assimilation and transcranial magnetic stimulation-evoked potentials in group 1. **a**) Correlations of excitatory gain parameter *A* and intracortical facilitation (ICF). **b**) Correlations of inhibitory gain parameter *B* and short-interval intracortical inhibition (SICI). Each panel shows scatter plots of the correlation of the transcranial magnetic stimulation-evoked potentials (ICF or SICI) with the E/I gain parameters (*A* or *B*). The *r*_*sp*_ and *p* values indicate the Spearman correlation coefficient and its *p*-value evaluated by Robust Correlation Toolbox [34], respectively. The *p*-values were corrected with the Bonferroni method to avoid the multiple comparison effect. The *r*_*sp*_ and *p* values in red indicate statistically significant correlations (*p <* 0.05 with Bonferroni correction).

**Fig. 5.**
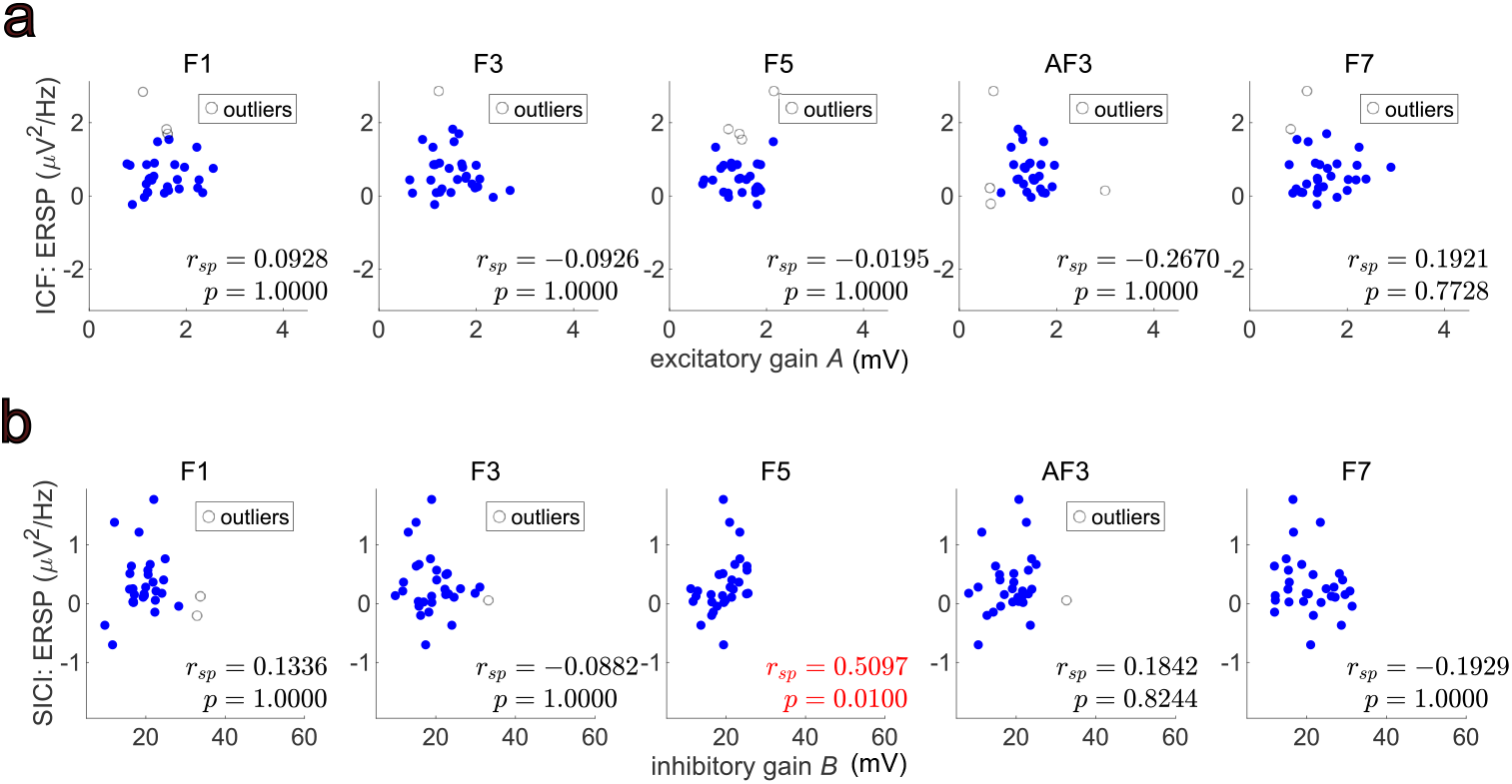
Excitation/inhibition (E/I) estimations based on data assimilation and event-related spectral perturbation (ERSP) in group 2. **a**) Correlation of excitatory gain parameter *A* and excitatory ERSP. **b**) Correlation of inhibitory gain parameter *B* and inhibitory ERSP. Each panel shows scatter plots of the correlations of the E/I ERSP with the E/I parameters. The *r*_*sp*_ and *p*-values indicate the Spearman’s correlation coefficient and its *p*-value evaluated by Robust Correlation Toolbox [34], respectively. The p values were corrected with the Bonferroni method to avoid the multiple comparison effect. The *r*_*sp*_ and *p* values in red indicate significant correlations (*p <* 0.05 with Bonferroni correction).

The estimated excitatory gain parameter *A* was compared with the ICF features indexed by P60 TEP components in group 1 (Fig. 4a). Moreover, the estimated inhibitory gain parameter *B* was compared with the SICI features indexed by N100 TEP components in group 1 (Fig. 4b). In the intracortical excitation and inhibition comparisons, the DA-based features indexed by the estimated parameters in the NM model were significantly correlated with the TEP features (Fig. 4); however, these significant correlations showed in the limited electrode (excitatory gain parameter *A* vs ICF - F5, *r*_*sp*_ = 0.5374, *p* = 0.0066 / inhibitory gain parameter *B* vs SICI -F3, *r*_*sp*_ = 0.4612, *p* = 0.0295). The other electrodes showed no significant correlations in either comparison (excitatory gain *A* vs ICF - F1, *r*_*sp*_ = 0.0167, *p* = 1.0000; F3, *r*_*sp*_ = 0.1961, *p* = 0.7701; AF3, *r*_*sp*_ = −0.0788, *p* = 1.0000; F7, *r*_*sp*_ = −0.2325, *p* = 1.0000 / inhibitory gain parameter *B* vs SICI - F1, *r*_*sp*_ = −0.1138, *p* = 1.0000; F5, *r*_*sp*_ = 0.3306, *p* = 0.1996; AF3, *r*_*sp*_ = −0.5000, *p* = 1.0000; F7, *r*_*sp*_ = 0.1200, *p* = 1.0000).

The correlation analysis results of DA-based E/I features (E/I gain parameters *A* and *B*) and ICF/SICI features indexed by ERSP of gamma-band in group 2 are shown in Fig. 5a, b. In contrast to the results of Fig. 4, even though the comparison of the inhibitory component (inhibitory gain parameter *B* vs SICI) showed a significant correlation in F5 (F1, *r*_*sp*_ = 0.1336, *p* = 1.0000; F3, *r*_*sp*_ = −0.0882, *p* = 1.0000; F5, *r*_*sp*_ = 0.5097, *p* = 0.0100; AF3, *r*_*sp*_ = 0.1842, *p* = 0.8244; F7, *r*_*sp*_ = −0.1929, *p* = 1.0000), that of the excitatory component (excitatory gain *A* vs ICF) showed no significant correlation for any of the electrodes (F1, *r*_*sp*_ = 0.0928, *p* = 1.0000; F3, *r*_*sp*_ = −0.0926, *p* = 1.0000; F5, *r*_*sp*_ = −0.0195, *p* = 1.0000; AF3, *r*_*sp*_ = −0.2670, *p* = 1.0000; F7, *r*_*sp*_ = 0.1921, *p* = 0.7728).

## Discussion

In this study, we directly compared E/I estimation results derived from the proposed DA-based method and neurophysiological indices (SICI/ICF) using the TMS-EEG method to validate our proposed method. The results showed that E/I balance in the prefrontal cortex was retained across participants with both methods (our proposed method and the TMS-EEG neurophysiological method). Furthermore, the E/I gain parameters *A* and *B*, estimated from the resting-state EEG only by our proposed method, showed significant correlations with the ICF and SICI components indexed by TEP, respectively. That is, we demonstrated that our proposed DA-based method can successfully assess changes in E/I balance without contradicting results based on TMS neurophysiological indices using the TMS-EEG method. To date, several data-driven NM modeling [37–39] and E/I estimation methods [40, 41] based on EEG have been proposed in human EEG studies. However, these methods still lack evidence to support the neurophysiological validity of the estimated model parameters (e.g., E/I synaptic gains and time constants of E/I interneuron columns). Therefore, the findings of this study suggest a remarkable advantage and neurophysiological validity of the estimated E/I parameters obtained from our proposed DA-based method. In the following sections, we provide detailed discussions regarding the neurophysiological validity and advantages of the DA-based E/I estimation method.

The concurrent TMS-EEG measurement technique [42, 43] has been developed as a non-invasive method to measure changes in E/I balance in the human cortex under both healthy and pathological conditions [3, 5, 17, 18]. As described in the Methods section, the TMS-EEG method enables the application of an appropriate paired-pulse TMS paradigm to detect the facilitatory (ICF) and inhibitory (SICI) functions in the targeted cerebral cortex. Practical knowledge and neurophysiological evidence accumulated over the past 15 years using TMS-EEG techniques are gradually unraveling the functional mechanisms of TMS-elicited EEG [44]. Several previous TMS-EEG studies have experimentally shown that TEP changes reflect both inhibitory and excitatory functions mediated by GABA and glutamate receptors, respectively [18, 29, 31]. For example, the N100 component of the TEP obtained in the SICI paradigm reflects inhibitory function mediated by GABA_A_ receptors [17, 30]. Conversely, the P60 component of the TEP obtained in the ICF paradigm is mainly associated with glutamatergic excitatory function via *α*-3-hydroxy-5-methyl-4-isoxazole propionic acid (AMPA) receptors and inhibitory function via GABA_A_ receptors [18, 30, 31].

Another method for measuring E/I balance is MRS. Recent studies suggest that MRS can measure E/I balance changes associated with perceptual and motor learning [13, 16, 45]; however, the neurophysiological validity of measuring E/I balance with MRS remains to be confirmed. MRS cannot discriminate between extrasynaptic and synaptic responses of GABA and glutamate [14], and thus total GABA concentrations obtained from MRS do not necessarily reflect the dynamics of GABA receptor-mediated neurophysiological functions [14–16]. Taken together with the practical evidence from TMS-EEG and MRS studies, and considering that the TMS-EEG method is a reliable and established approach for measuring E/I balance, we examined the neuroelectrophysiological validity of our proposed method by directly comparing the ICF/ISCI indexed by the TMS-evoked EEG with the proposed DA-based E/I estimation method.

Comparing DA-based E/I gain parameters with ICF/SICI TEPs revealed significant correlations for both prefrontal excitation (the parameter *A* and ICF-TEP) and inhibition (the parameter *B* and SICI-TEP) in the L-DLPFC (i.e., the brain site of TMS) (see F3 in Fig. 4a and F5 in Fig. 4b). Moreover, in the DA-based E/I estimation analysis for resting-state EEG data, we applied artifact rejection based on independent component analysis [46] and current source density transformation [47, 48] as the preprocessing of EEG for the DA-based method. This improved spatial resolution relative to raw EEG. Therefore, these significant correlations in the limited electrodes between estimated parameters (*A* and *B*) and ICF/SICI-TEP amplitudes could be considered reasonable and acceptable from a neurophysiological viewpoint.

Conversely, ICF/SICI components indexed by ERSP of gamma-band were only correlated with the SICI features (the parameter *B* in F5 vs. ERSP of gamma-band). The lacking significant correlation between the excitatory gain parameter *A* and ERSP of gamma-band may be explained by the fact that GABA_A_ receptor-mediated inhibitory activity is more dominant (as the origin of brain regions for gamma oscillations) than AMPA glutamate receptor-mediated excitation [49]. Therefore, the gamma excitatory ERSP in the ICF TMS-EEG paradigm is masked by other background neuronal activity and observational noise. We believe that this masked effect of AMPA glutamate receptor-mediated excitation in ERSP of gamma-band led to a weak correlation with excitatory gain parameter *A*. Considering all of the above, the results indicate that our proposed DA-based method can estimate the E/I gain parameters reflecting neurophysiologically valid neural features regarding glutamate-mediated excitation and GABA-mediated inhibition.

Our EEG-DA algorithm originally proposed in [12] was modified in this study by combination with ETKF [28] (see the Methods section). As shown in the Supplementary Results regarding numerical validation with synthetic EEG data, introducing ETKF drastically decreased the number of ensemble members *N*_*ens*_ for state estimation with Sequential Monte Carlo sampling.

Although a requirement for accurate state estimation of Sequential Monte Carlo in the previous EnKF-based method (i.e., vbcEnKF [12]) was *N*_*ens*_ ≥ 200, the current ETKF-based method (i.e., vbcETKF) can achieve equal or more accurate estimation performance with only *N*_*ens*_ = 70 (see Supplementary Results for more details). As this achievement indicates a significant decrease in the computational cost for the state estimation of the NM model, our proposed DA-based data-driven E/I estimation method has the great advantage of lower costs. Moreover, our results indicate that the proposed DA-based method can estimate neurophysiologically valid E/I balance changes. Even though this study does not focus on the time-varying changes in E/I activity, our proposed method can track the time-varying changes in E/I balance at the sub-second scale because of its recursive Bayesian filtering scheme, as demonstrated in a previous study [12]. This is the most significant advantage of our proposed DA-based E/I estimation method compared with other conventional TMS-EEG-based E/I estimation methods. The TMS-EEG-based E/I estimation method requires analyzing the TMS-triggered averaged EEG and performing multiple (≥ 60) trials to identify and extract TEP components from TMS-evoked EEGs [50, 51]. By contrast, our method has the advantage of detecting the moment-to-moment changes in E/I balance from a short-term resting-state EEG. Taken together, our method could be applied to advance our understanding of neuroplasticity. Furthermore, the neurophysiological validity of our proposed method in this study indicates that the method is not only applicable to basic neuroscience research to elucidate the functional mechanisms underlying changes in E/I balance, but may also elucidate the pathophysiology of neuropsychiatric disorders based on E/I balance and thus facilitate the development of clinical treatments. For example, our method can be applied to develop a neurofeedback system as indexed by the E/I balance changes in sub-second order. It would be helpful to improve the strategy of neuromodulation-based rehabilitation for patients with neuropsychiatric disorders by combining neurofeedback based on TMS neuromodulation and E/I balance. We believe that such a combined method will contribute to the establishment of novel strategies for E/I state-informed neuromodulation.

This study further validated and improved the computational efficiency of our previously proposed data-driven method for estimating changes in E/I balance that was based on the NM model using the DA scheme [12].

Although this study confirmed the advantages of our modified method, some limitations still exist. First, our proposed method retains the effects of hyper-parameter settings. For instance, the scale of state noise covariance *Q* is fixed in our proposed method. Even though the observation noise covariance *R* is adaptively estimated from observed data using the variational Bayesian noise-adaptive algorithm for state-space modeling [20–23], the state noise covariance *Q* is still fixed in our method. One of the simple solutions to this issue is modifying our method based on the self-organizing KF algorithm proposed previously [52]. The self-organizing KF can estimate both noise covariance *Q* and *R* in parallel with state and parameter estimation by adding the covariance *Q* and *R* for target-to-state estimation. However, to address these issues, further studies are required. Second, whether our method can generally track the E/I ratio changes during cognitive tasks is still unclear. Based on evidence from our previous [12] and current studies, we confirmed two important findings: (1) our method can detect sleep-dependent E/I balance changes, and (2) the estimated E/I gain parameters *A* and *B* are neurophysiologically valid. However, to reveal whether our method can estimate the transient E/I balance changes during general tasks, we should apply our proposed method to the EEG dataset obtained from participants while conducting cognitive tasks (e.g., cognitive and motor learning tasks). Moreover, in future studies, we will modify our DA-based method to enable its application to estimations of network connectivity in parallel with tracking E/I balance changes.

## Methods

### Participants

Healthy participants were screened and recruited at Keio University Hospital between 2017 and 2022. In total, 29 participants (mean age = 45.9 ± 13.3 years; 18 men, 11 women) were included in the TEP analysis, and 30 participants (mean age = 33.4 ± 11.1 years; 14 men, 16 women) were included in the time-frequency analysis. The screening was performed by certified psychiatrists to confirm the absence of a history of psychiatric disorders through the structured clinical interview for DSM (Diagnostic and Statistical Manual of Mental Disorders) [53]. The study protocol (UMIN000028863) was approved by the Ethics Committee of the Keio University School of Medicine. All participants provided written informed consent in accordance with the Declaration of Helsinki.

### Data collection: concurrent TMS-EEG recording

For the TMS experiments, we used a monophasic TMS stimulator (DuoMAG MP stimulator; DEYMED Diagnostic Ltd., Hronov, Czech Republic) equipped with a 70 mm diameter figure-8 butterfly coil (DuoMAG 70BF; DEYMED Diagnostic Ltd.). We determined coil location with online monitoring by importing T1-weighted images into the Brainsight TMS Navigation system (Rogue Research Inc., Montréal, QC, Canada) and registering them with digitized anatomical landmarks. EEG was recorded using a TMS-compatible 64-channel amplifier (TruScan LT, DEYMED Diagnostic s.r.o., Hronov, Czech Republic) equipped with a sample-and-hold circuit. For the EEG cap, silver C-ring slit electrodes were used (TruScan Research EEG Caps, 64-channel, DEYMED Diagnostic s.r.o., Hronov, Czech Republic). The sampling rate of the EEG data was 3 kHz. The reference electrode was placed on the right earlobe and the ground electrode on the left earlobe. The resting motor threshold (RMT) was measured based on muscle activity in the first dorsal interosseous muscle of the right hand using electromyography after identifying the optimal stimulation site on the left primary motor cortex. RMT was defined as the minimum intensity that elicited motor-evoked potentials of ≥ 50*µV* in the target muscle in half of the TMS trials to the left primary motor cortex with the EEG cap on.

Following previously established methods [31], single-pulse and paired-pulse TMS were performed on the left DLPFC of each participant at 5 s (±0.5 s) intervals, at inter-stimulus intervals of 2 ms for SICI and 10 ms for ICF. Furthermore, 80 trials of single-pulse TMS (test pulse at 120% of RMT intensity) and 80 trials of paired-pulse TMS (condition pulse at 80% of RMT intensity + test pulse at 120% of RMT intensity) were randomly administered to each participant.

During TMS stimulation, the coil was placed at a 45 degree angle to the midline and the stimulation sites were individually placed at the MNI coordinates [*x* = −38, *y* = 44, *z* = 26] using the MRI-guided neuronavigation system (Brainsight).

Additionally, we applied a white noise masking method to all participants using an earplug-type sound stimulator to suppress auditory-evoked potentials elicited by TMS click sounds. The volume was individually adjusted to cancel out the TMS click sounds [54], and foam was placed directly under the coil for TMS-EEG measurements.

### Data collection: resting EEG recording

EEG was recorded using a TMS-compatible 64-channel amplifier (TruScan LT, DEYMED Diagnostic s.r.o., Hronov, Czech Republic) equipped with a sample-and-hold circuit. Resting-state EEG was measured in a relaxed awake state with eyes closed for approximately 5 min.

### Preprocessing of EEGs

Preprocessing of the TMS-EEG data was conducted using EEGLAB v2021.0 [55], TMS-EEG Signal Analyzer (TESA v1.1.1) [56], and custom scripts in MATLAB (R2020a, the MathWorks Inc., Natick, MA, USA).

For resting-state EEG analysis, EEG data were resampled to 250 Hz, and high-pass and low-pass filters (1–50 Hz) were applied. Additionally, 50 Hz line noise was removed using the cleanline function [57]. Noisy channels and time segments contaminated with extreme noise were automatically identified and removed using the validated Artifact Subspace Reconstruction algorithm [58]. Each subject’s EEG data was re-referenced, and then independent component analysis was performed. After independent component analysis decomposition, the ICLabel algorithm (a semi-automatic method for component selection) was applied [59]. For TMS-EEG analysis, EEG data was initially segmented between −2000 ms and 2000 ms. Baseline correction was performed by subtracting the average signal amplitude between −500 ms and −150 ms. Electrodes exhibiting high variability (median z-scores *>* 3) were automatically excluded. Epochs with excessive noise (*>* 1000*µV*) were also automatically removed, and remaining noisy epochs were manually inspected and eliminated. Specific electrodes (F5, F3, F1, F7, AF3, FC3, and FC5) corresponding to the DLPFC stimulation site were exempt from automatic exclusion. To avoid TMS pulse artifacts, EEG data from −5 ms to 25 ms was removed. Cubic interpolation was applied to prevent ringing artifacts during downsampling and filtering. The data was downsampled to 1 kHz and underwent an initial round of fast independent component analysis to eliminate physical TMS decay components. Subsequently, the data was filtered using bandpass (0.5–100 Hz) and notch (48–52 Hz) filters. Removed channels underwent spherical interpolation. A second round of independent component analysis (EEGLAB infomax (runica)) was applied to remove artifacts, such as eye blinks, eye movements, and muscle artifacts. Finally, the data was re-referenced to the overall electrodes.

### EEG analysis for concurrent TMS-EEG recording

TMS-EEG analysis was performed at the sensor level, with the F3, F5, and AF3 electrodes corresponding to the left DLPFC. The TEP was calculated as the electrode average of the total trial average at each electrode. Then, SICI-TEP and ICF-TEP were calculated as the difference between the TEP obtained from the SICI or ICF paradigms and the TEP obtained from single-pulse TMS. Specifically, SICI-TEP was calculated using the negative peak (N100) of the TEP within the 70–130 ms range, induced by SICI and single-pulse stimulation. Similarly, ICF-TEP was calculated using the positive peak (P60) of the TEP within the 30–80 ms range, induced by ICF and single-pulse stimulation.

To investigate the oscillatory characteristics elicited by SICI and ICF paradigms applied to calculate ERSP for single-pulse TMS, SICI, and ICF. ERSP represents the time-locked spectral activity induced by TMS relative to a prestimulus baseline. ERSP was computed by averaging power over epochs at specified channel-frequency time points using open-source minimum norm estimate software (tfr morlet) with Morlet wavelets ranging from 3 to 10 cycles [32]. A frequency range of 4–48 Hz and time interval of 30–250 ms were included in the analysis. ERSP was baseline-corrected (−500 to −100 ms) and converted to logarithmic ratios. Additionally, we calculated the mean values in the gamma band (30–48 Hz) and the 40–60 ms time range at the left DLPFC stimulation site (average of the F1, F3, F5, F7, and AF3 electrode sites). Then, SICI-ERSP was calculated by subtracting the mean ERSP obtained from the SICI paradigm from the mean ERSP obtained from single-pulse TMS. Similarly, ICF-ERSP was calculated by subtracting the mean ERSP obtained from single-pulse TMS from the mean ERSP obtained from the ICF paradigm.

### Descriptions of E/I balance tracking based on the NM model

Here, we describe the DA-based evaluation of E/I balance from observed EEGs obtained in a resting condition. Before giving the mathematical details of our proposed EEG-DA method and E/I balance evaluation, we will first briefly describe our proposed method. In our method, the NM model is directly assimilated with the observed EEG, and E/I balance changes are tracked by calculating the time-varying ratio of the estimated parameters regarding E/I synaptic gains [12]. The NM model, proposed by Jansen and Rit [60], is well known as a computational model for EEG signals that is formulated as the interaction between three neural cell populations: pyramidal cells, excitatory interneurons, and inhibitory interneurons. This model can be described as the following six first-order differential equations [60]:

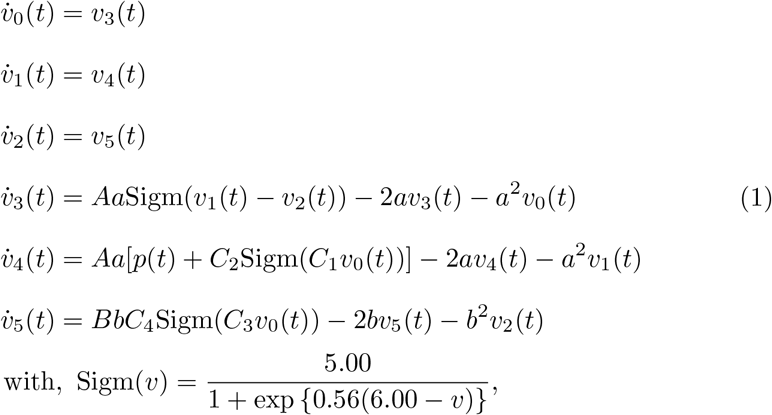

where variables *v*_0_, *v*_1_, and *v*_2_ indicate the mass of postsynaptic potential (PSP) in three neural populations: pyramidal cells, excitatory interneurons, and inhibitory interneurons, respectively. The synthetic EEGs generated in the NM model are defined as *y*(*t*) = *v*_1_(*t*) − *v*_2_(*t*). The fixed constants *C*_*n*_ (where, *n* = 1, …, 4) account for the number of synapses established between two neural populations. *C*_*n*_ was fixed as *C*_1_ = 135, *C*_2_ = 108, *C*_3_ and *C*_4_ = 33.75, based on a previous study [61]. The other five parameters in the NM model, *A, a, B, b*, and *p*, are the target-to-state estimation in our proposed DA method. Parameters *A* and *B* represent the excitatory and inhibitory synaptic gains, which control the amplitude of the excitatory PSP (EPSP) and inhibitory PSP (IPSP), respectively. Parameters *a* and *b* indicate the inverse of the time constant for EPSP and IPSP that control the dominant frequency band of the generated EEG oscillations in the NM model. Parameter *p* represents the background noise input of the model.

To apply our proposed EEG-DA method, the six first-order differential equations in the model were transformed to the state-space form as follows:

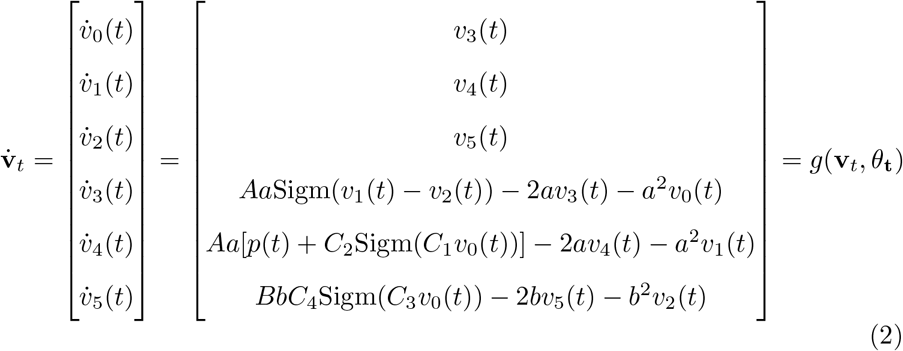

state model;

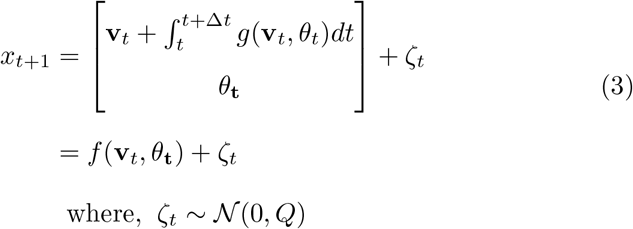

observation model;

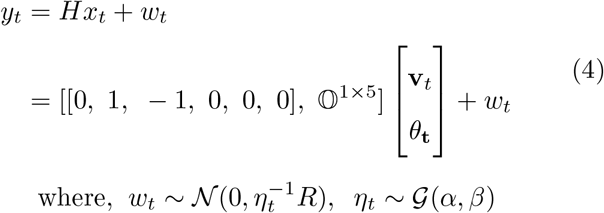

where *v*_*t*_ = {*v*_0_(*t*), *v*_1_(*t*), …., *v*_5_(*t*)} and *θ* = {*A, a, B, b, p*} indicate the model variables and parameters of the NM model, respectively. Note that 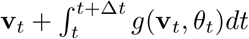 was implemented with the Euler method with step size Δ*t* (sampling interval). The state *x*_*t*_ = {*v*_*t*_, *θ*_*t*_}^*T*^ and observation *y*_*t*_ follow Gaussian distributions as *x*_*t*_ ∼ 𝒩 (*x*_*t*_, *P*_*t*_) and 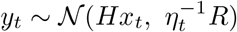,respectively. The variable *ξ*_*t*_ is state noise that follows the normal density *ξ*_*t*_ ∼ 𝒩 (0, *Q*). Variable *w*_*t*_ is observation noise, which follows the normal distribution 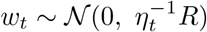 To consider the nonstationary noise effect, we introduced the noise scale factor *η*_*t*_ that follows the gamma probability *η*_*t*_ ∼ 𝒢 (*α, β*). In this case, the state *x*_*t*_ and noise scaling factor *η*_*t*_ can be estimated from the observation *y*_*t*_ by using the following Bayesian theory:

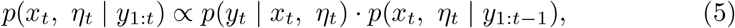

where *y*_1:*t*_ indicates the observation set as {*y*_1_, *y*_2_, …, *y*_*t*_}. If *x*_*t*_ and *η*_*t*_ can be defined as independent, the posterior distribution *p*(*x*_*t*_, *η*_*t*_ | *y*_1:*t*_) in equation 5 can be solved by minimizing the Kullback–Leibler (KL) divergence between the approximated distribution *q*(*x*_*t*_)*q*(*η*_*t*_) and exact distribution *p*(*x*_*t*_, *η*_*t*_ | *y*_1:*t*_) [20]:

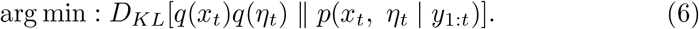

In our proposed method, by combining equation 6[12, 20–23] with the ETKF [28], we can derive solution of the state *x*_*t*_ = {*v*_*t*_, *θ*_*t*_}^*T*^ and noise scaling factor *η*_*t*_. The detailed description for the solution of equation 6 is described in the following section.

Based on this Bayesian estimation scheme, the time-varying changes in the model variable *v*_*t*_ and parameter *θ*_*t*_ can be estimated from the observed EEG *y*_*t*_ on a sample-by-sample basis while also considering nonstationary noise effects. As the estimated parameters *θ*_*t*_ contained the parameters representing the E and I synaptic gains *A*(*t*) and *B*(*t*) in the NM model, time-varying changes in E/I balance can be quantified by calculating the ratio of these parameters. Thus, we defined the model-based evaluation index of the E/I ratio as mE/I ratio:*mE/I*(*t*) = *A*(*t*)*/*(*A*(*t*) + *B*(*t*)), which can be calculated from the estimated parameters *θ*_*t*_ = {*A, a, B, b, p*}.

### Variational Bayesian noise-adaptive EnKF

In the following sections, we will describe the solution of equation 6 and provide details of our proposed EEG-DA algorithm. First, we will explain the solution of equation 6 in a linear Gaussian state-space model. In this case, the state-space model is formulated with the following linear equations:

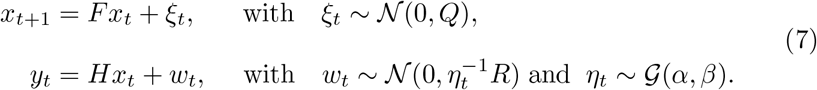

Here, we assumed that the probability of the observation noise follows the normal gamma probability density. Furthermore, observation *y*_*t*_ and state variable *x*_*t*_ follow the Gaussian distribution 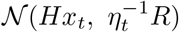 and 𝒢 (*x*_*t*_, *P*_*t*_), respectively. In this case, the variational Bayesian estimation as in equation 6 can be applied to estimate the state *x*_*t*_ and noise scaling factor *η* in the equation 7 [20–23]. By minimizing the KL divergence in equation 6 with respect to the probability *q*(*x*_*t*_), the solution of updating rule of the probability density for *x*_*t*_ is given as the following equation:

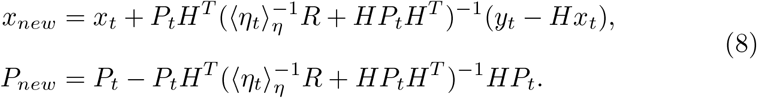

Note that ⟨·⟩_*η*_ indicates the expectation operator under the variable *η*. In the same manner, the solution of *q*(*η*) is also given, as follows:

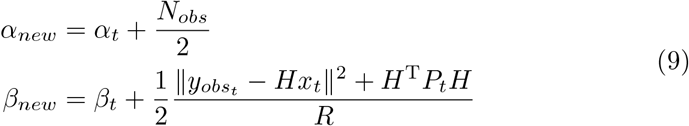

The expectation of noise scaling factor *η* (i.e., ⟨*η*⟩_*η*_) can be described as follows:

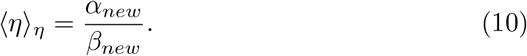

Based on equations 7–10, the procedures of the estimation in the linear Gaussian state-space model with the variational Bayesian noise-adaptive algorithm (vbKF) [20, 21] are as follows:

#### Algorithm 1

The variational Bayesian noise adaptive linear Kalman Filter (vbKF) algorithm

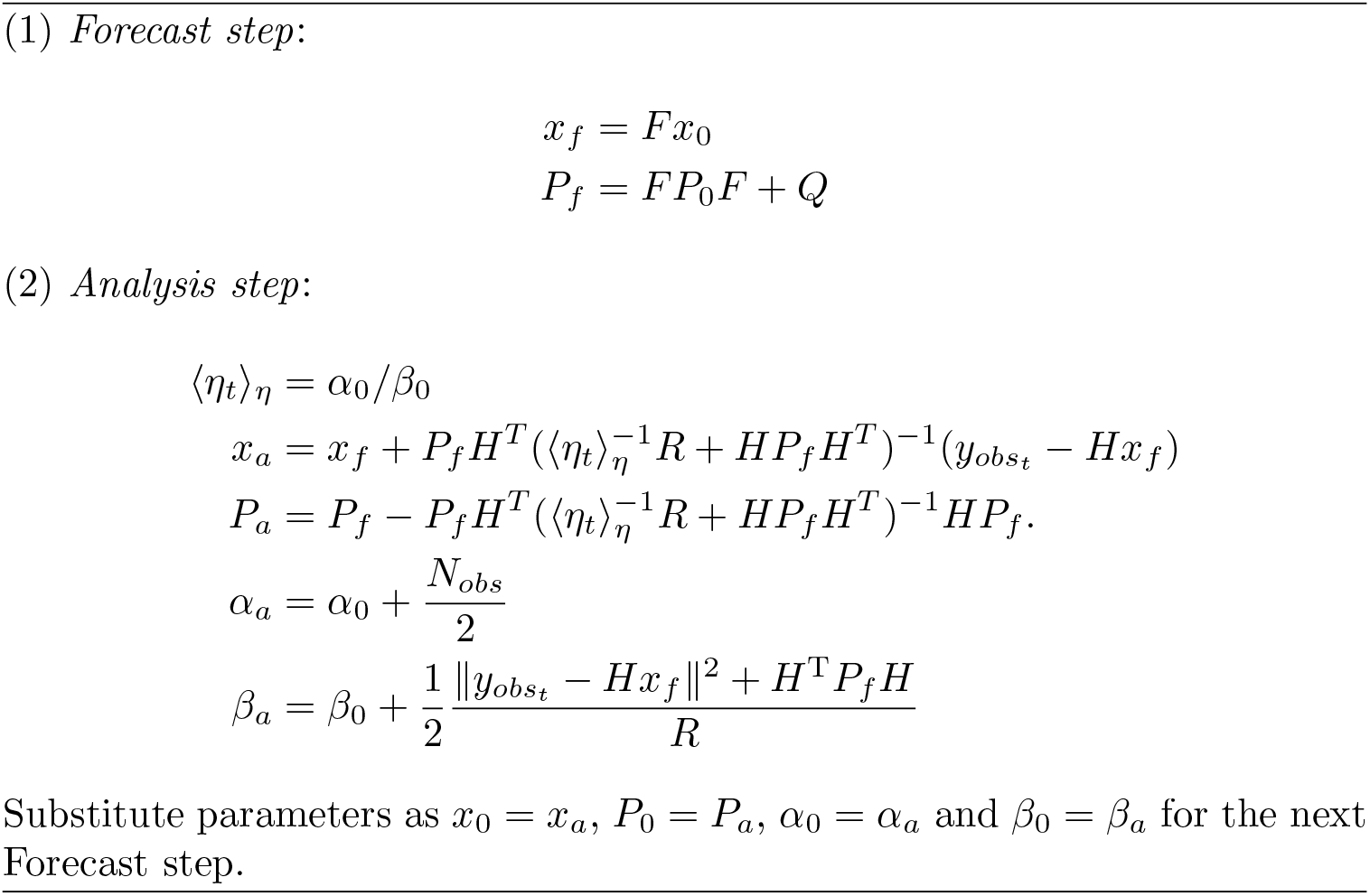

However, the linear vbKF approaches cannot directly be applied to the EEG-DA task because the time-evolving dynamics in the observed EEGs would be formulated as nonlinear dynamical systems, such as the NM model [60]. A simple solution to adapt vbKF approaches to nonlinear cases is combining the EnKF scheme with vbKF. In our previous work, we applied a combined method using the above noise-adaptive algorithms and EnKF (called vbEnKF) [12]. By using vbEnKF, algorithm 1 can be extended for the estimation of the nonlinear state-space model as shown below:

#### Algorithm 2

The variational Bayesian noise-adaptive EnKF (vbEnKF) algorithm

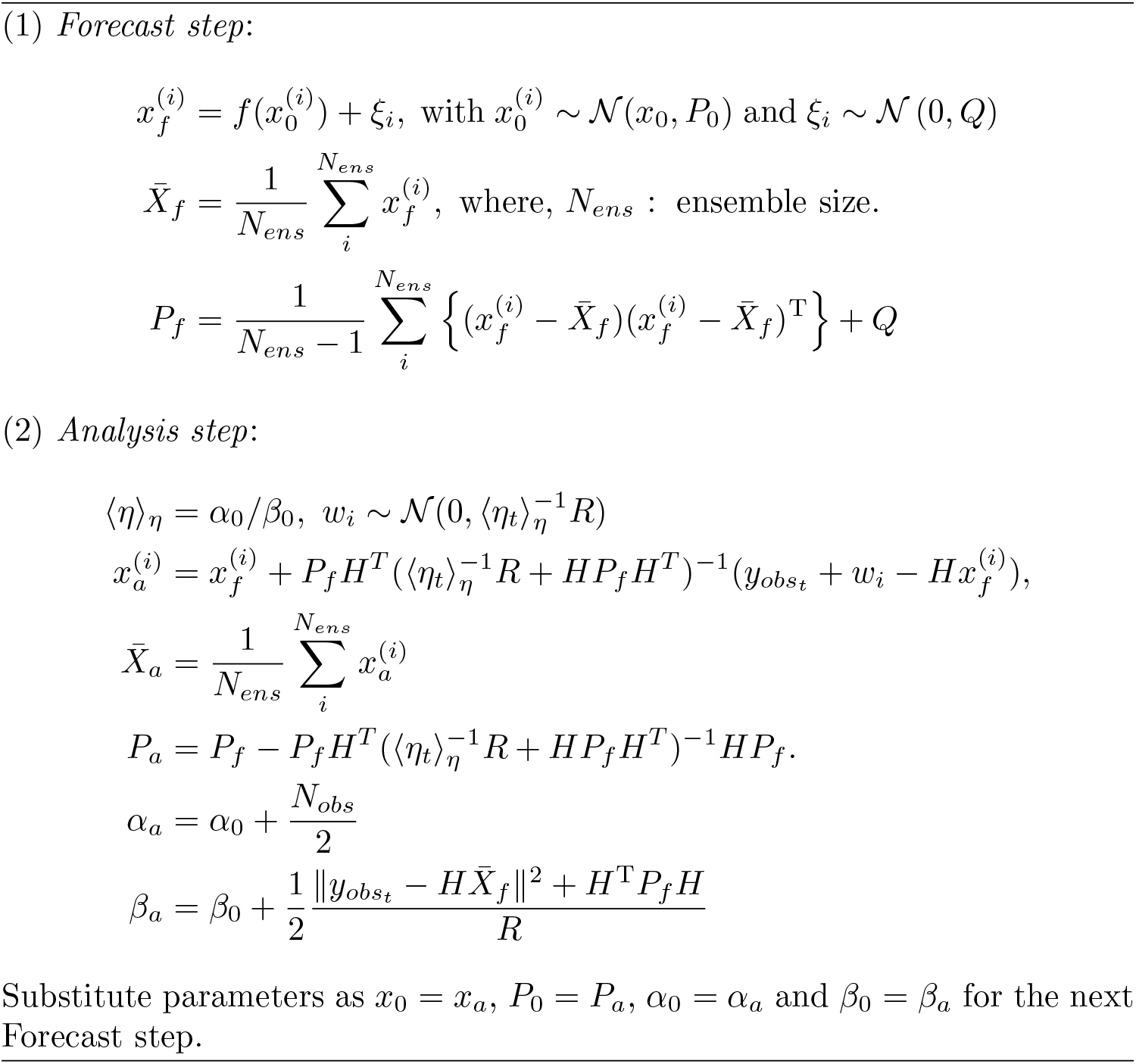

### The constrained Kalman Filter

We also applied the constrained Kalman Filter scheme for state estimation in the NM model to consider the boundary constraint of parameter *θ*_*t*_. If state *x*_*t*_ follows the probability density *x*_*t*_ ∼ *p*(*x*_*t*_ | *x*_*t*−*t*_) = 𝒢 N(*x*_*t*_, *P*_*t*_) with the constraint *d*_*L*_ ≤ *Dx*_*t*_ ≤ *d*_*U*_, the algorithm estimates the value of *x*_*t*_ and *P*_*t*_ so as to solve the following problem:

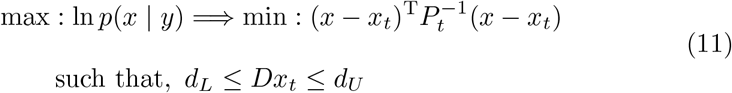

This problem can be solved using a Lagrange multiplier method and the solution is given as follows [24–27]:

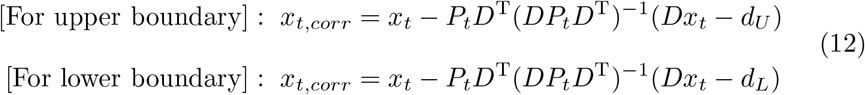

Equation 12 is applied only when the estimation of *x*_*t*_ based on the solution of equation 6 violates the boundary constraints.

### New proposed method: variational Bayesian noise-adaptive constrained Ensemble Transform Kalman Filter (vbcETKF)

In our previous study [12], by combining vbEnKF and the constrained KF scheme, we proposed the NM model-based EEG-DA method using the vbcEnKF scheme [12] to consider both nonstationary noise effects and parameter constraints. EnKF is well known as the DA algorithm for predicting time series of nonlinear dynamical systems. However, the vbcEnKF scheme is used for the Perturbed Observation (PO) method-based EnKF (PO-EnKF) [35, 62] scheme, and thus this method is required for large numbers of ensembles for estimating the probability density of state variables. This critically affects the computational cost (e.g., processing time and memory requirements) for DA. In the current study, to reduce the costs, we modified the vbcEnKF scheme by combining it with the ETKF [28] scheme. In this section, we will describe the basics of ETKF and how to apply ETKF for our proposed EEG-DA method.

PO-EnKF is required to add the randomized perturbation*w*_*i*_ ∼ 𝒢 (0, *R*) for the observations to estimate the analyzed state *x*_*a*_ as in algorithm 2, because observations without treating a perturbing noise led to underestimation of state covariance and filter divergence [35, 36]. However, such treatment of perturbing noise for observations increased the number of ensemble members for the state estimation with Sequential Monte Carlo sampling. To solve this issue, Bishop and colleagues [28] proposed the ETKF to establish ensemble filtering without perturbed observations. In this method, instead of treading the perturbed noise *w*_*i*_ in observations, Bishop and colleagues [28] supposed that analysis error of state 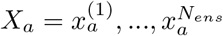 can be explained as state perturbation 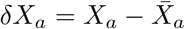 with 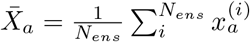.Moreover, by introducing the linear operator *T*, the authors assume that analyzed error of state *δx*_*a*_ can explain linear transformation of forecasting error of state *δx*_*f*_ by *T* .

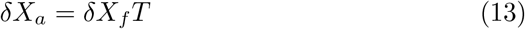

where 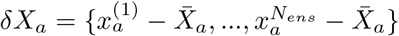 and 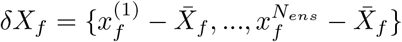. Based on equation 13, the update rule of state *X*_*a*_ in the analysis step of the standard PO-EnKF [35, 62] scheme can be rewritten as below.

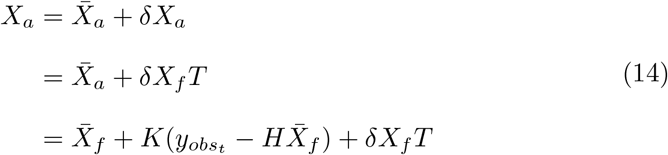

where, *K* indicates the Kalman gain: *K* = *P*_*f*_ *H*^*T*^ (*R* + *HP*_*f*_ *H*^*T*^)^−1^. Moreover, the analyzed state covariance *P*_*a*_ should satisfy the following equation.

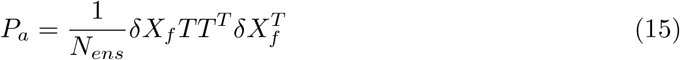

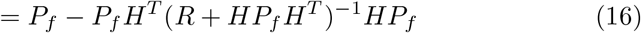

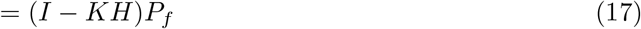

To solve linear operator *T* so as to satisfy equation 15 = equation 17, the Kalman gain *K* and forecasting state covariance *P*_*f*_ in equation 17 are written as follows:

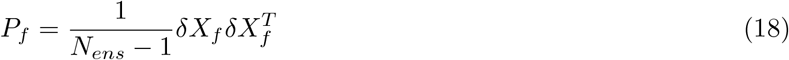

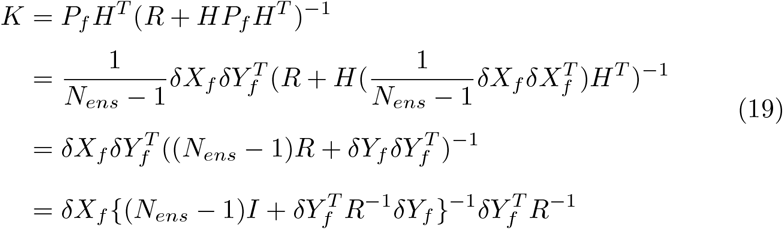

where, 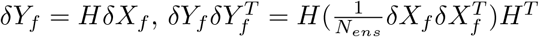 and 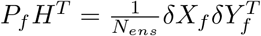.

By substituting equations 18 and 19 into equation 17, the analyzed state covariance *P*_*a*_ can be rewritten as follows:

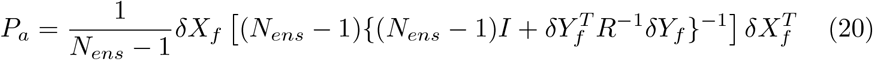

From equation 15 = equation 20, the linear operator *T* can be solved as follows:

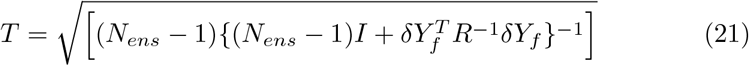

Finally, by using equations 19, the update rule of state *X*_*a*_ in the analysis step as in equation 14 can be rewritten as follows:

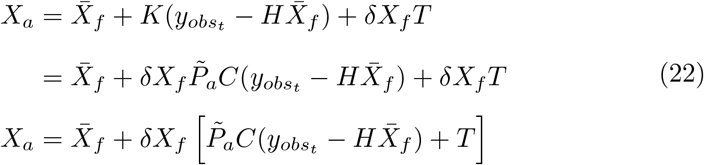

where 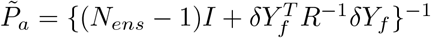 and 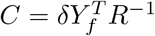.

Based on the above, our proposed algorithms for EEG-DA are modified by combining the previously proposed vbcEnKF scheme [12] with ETKF [28]. The entire algorithm of the modified EEG-DA method (called vbcETKF) is summarized in Algorithm 3.

#### Algorithm 3

The variational Bayesian noise adaptive constrained Ensemble Transform Kalman Filter (vbcETKF) algorithm

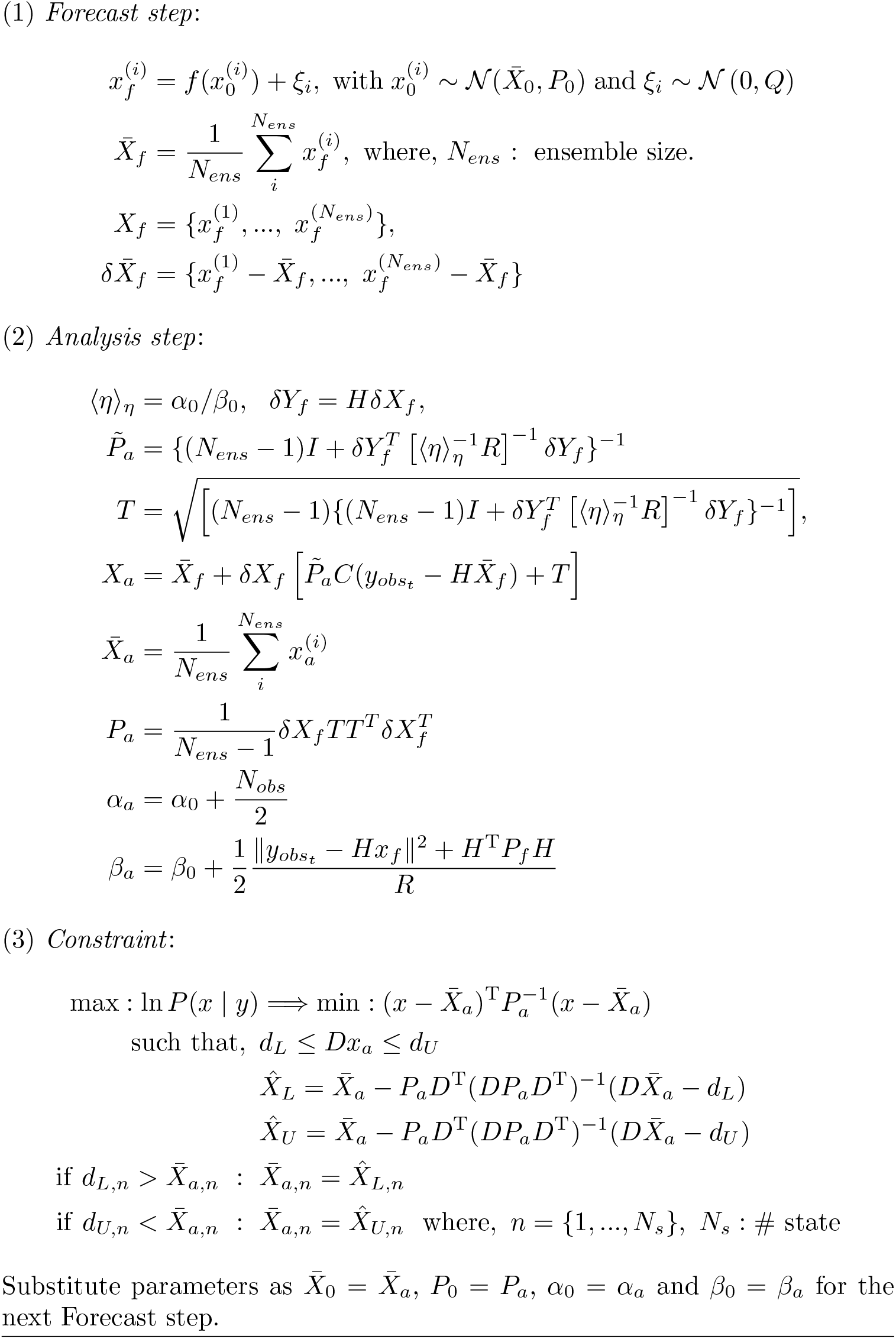

By applying the above-mentioned state-space form of the NM model to the vbcETKF algorithm, the model state *v*_*t*_ = {*v*_0_, *v*_1_, …, *v*_5_} and parameters *θ*_*t*_ = {*A*_*t*_, *a*_*t*_, *B*_*t*_, *b*_*t*_, *p*_*t*_} were recursively estimated from observed EEG data on a sample-by-sample basis. The entire process of this method includes the following steps:

1. Initialize the parameters for the probabilities 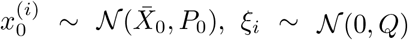, and *η*_0_ ∼ 𝒢 (*α*_0_, *β*_0_).
2. Calculate the one-step ahead prediction of state ensemble members 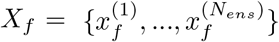 and its average 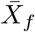 (see the Forecast step in Algorithm 3)
3. Calculate the analyzed parameters in state variables (*X*_*a*_ and *P*_*a*_) and observational noise scale factor (*α*_*a*_ and *β*_*a*_) (see the Analysis step in Algorithm 3)
4. If the mean of state variables 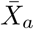 violate the constraint, the corrected state value is calculated so that 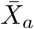 is within a feasible range (see the Constraint step in Algorithm 3).
5. Substitute all updated parameters as 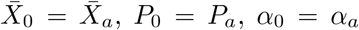 and *β*_0_ = *β*_*a*_ for the next Forecast step.
6. Repeat steps 2–5.

For step 1, the initial state variable 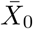 was set at 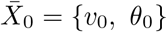,where *v*_0_ = **O**^1*×*6^,and *θ*_0_ = {*A*_0_, *a*_0_, *B*_0_, *b*_0_, *p*_0_}. Note that the initial value of NM model parameter *θ*_0_ is set as the standard parameter setting to produce alpha-like EEG oscillations from previous study [60]. The observation noise covariance R was set at *R* = 50. The initial setting for G(*α*_0_, *β*_0_) was set at *α*_0_ = 10^−3^ and *β*_0_ = 10^−3^. The state noise covariance *Q* was *Q* = diag ([Δ*t* · 𝕀 ^1*×*6^, 10^−3^ · 𝕀 _1*×*5_]).

In the constraint step of Algorithm 3, we applied the following interval constraint:

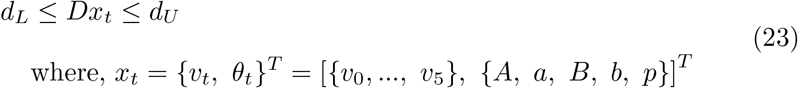

The following parameters were set: *D* = [𝒪 ^5*×*6^, **I**^5*×*5^], *d*_*L*_ = [0.01, 5.00, 0.01, 5.00, 120]^*T*^, and *d*_*U*_ = [100.00, 200.00, 100.00, 200.00, 320.00]^*T*^ . Such interval constraints were set to restrict the range of values in the model parameters *A, a, B, b*,and *p*, drawing from our previous work [12].

As mentioned above, the E/I balance is evaluated as the temporal changes in the ratio of the E/I gain parameters in the NM model (*A* and *B*), which is estimated from observed EEGs by using the vbcETKF scheme.

Therefore, we proposed the following calculation of the model-based index of the E/I ratio (mE/I ratio):

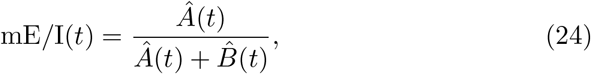

where 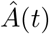 and 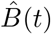 are putative E and I synaptic gain parameters based on our proposed vbcETKF method (Algorithm 3).

### EEG signal processing

After artifact removal, resting-state EEGs underwent current source density transformation [47, 48]. These transformed EEGs were bandpass filtered between 1 and 20 Hz with a zero-phase finite impulse response filter (number of taps = 3000; transition with 0.1 Hz; window function = Hanning window). Then the above-mentioned five electrodes located around the left DLPFC were selected, and the EEGs were individually assimilated to the NM model by using our proposed vbcETKF-based DA method to assess the time-varying E/I parameters.

### Comparison of E/I estimations based on DA and TEP

To compare the E/I estimation of our proposed method with the TMS-EEG-based method, we compared the E and I gains parameters (*A* and *B*) obtained from resting-state EEG with excitatory and inhibitory TEP components using correlation analysis. DA-based E/I were accessed by temporally averaged *A* and *B* gain parameters during a 5-minute resting condition. The E and I gain parameters were evaluated in the five electrodes located around L-DLPFC. TEP values were also calculated as the averaged value of the electrodes around L-DLPFC with two methods based on either conventional TEP components (N100 and P60) or ERSP (TMS-evoked EPSP in gamma-band 30–48 Hz).

The correlation analyses between DA-based and TEP-based E/I values were conducted using the Spearman’s rank correlation method with a box-plot-rule-based outlier removal algorithm in an open-source robust correlation MATLAB toolbox [34]. The *p*-value of the correlation coefficients were corrected with the Bonferroni method to correct for multiple comparisons.

## Acknowledgments

This research was partially funded by the National Institutes of Natural Sciences (NINS) program of Promoting Research by Networking among Institutions (grant number 01412303) and the NINS OPEN MIX LAB Program (grant number OML032401), the Japan Society for the Promotion of Science, Grant in Aid for Early-Career Scientists (grant number 18K15375), and Cooperative Study Program (25NIPS210) of the National Institute for Physiological Sciences.

## Data availability

Data will be made available to all interested researchers upon request.

## Code availability

Our implemented code of EEG-data assimilation method (vbcETKF) and its sample code of the numerical simulation described in Supplementary Methods are available at Github: https://github.com/myGit-YokoyamaHiroshi/EEGvbcETKFmEI_est.

## Author contributions

**Hiroshi Yokoyama**: Conceptualization, Formal analysis, Investigation, Methodology, Resources, Software, Validation, Visualization, Writing original draft, Writing – review & editing. **Yoshihiro Noda**: Funding acquisition, Investigation, Methodology, Supervision, Resources, Validation, Writing – review & editing. **Masataka Wada**: Data curation, Formal analysis, Investigation, Methodology, Software, Validation, Writing – review & editing. **Mayuko Takano**: Data curation, Formal analysis, Investigation, Methodology, Validation, Writing – review & editing. **Keiichi Kitajo**: Conceptualization, Funding acquisition, Project administration, Supervision, Resources, Validation, Writing – review & editing.

