## Supplementary information for "Validation of EEG data assimilation-based prefrontal excitation-inhibition balance estimation using TMS–EEG"

### Supplementary Results

#### S1. Numerical validation of our proposed method

*S1.1 Summary of our proposed method*

In the main manuscript, we proposed a modified version of our data assimilation (DA) method proposed previously [1] to estimate the changes in synaptic balance of excitatory and inhibitory neurons (E/I balance) from observed electroencephalographic signals (EEGs). This method can estimate E/I balance changes over time based on the observed EEGs while decreasing the computational cost relative to the previous DA method [1]. In this modified method, the observed EEGs obtained from a single electrode were directly assimilated into the NM model to estimate its model state and parameters based on the variational Bayesian constrained ensemble transform Kalman filter (vbcETKF). The vbcETKF is a direct extension of the previous version of our DA method called the variational Bayesian ensemble Kalman filter (vbcEnKF). As mentioned in the Methods section in the main manuscript, we successfully estimated E/I balance changes only from EEGs based on the vbcEnKF in our previous study [1]. However, EnKF is required to estimate the approximate solution of the NM model with a large number of ensembles with the sequential Monte Carlo sampling algorithm. Therefore, the ensemble size of sequential Monte Carlo sampling in the EnKF scheme directly affects the computational cost [2, 3]. To decrease such computational costs and enhance estimation efficiency, we proposed a modified version of our DA method using ETKF-based algorithms [4] instead of EnKF. As a result, by using ETKF, we confirmed that this modification can achieve a > 50% decrease in ensemble size for the state and parameter estimation in the NM model with numerical simulations (see section *S1.2*). In the following section, we describe the numerical validation of our proposed DA method in detail. Our findings reveal that vbcETKF exhibits enhanced computational efficiency compared with the previous method, vbcEnKF.

*S1.2 Simulation settings and results*

The main aim of this section is to reveal the numerical validity of our new proposed method, vbcETKF by comparing its estimation accuracy with that of the previously used vbcEnKF. In this numerical validation, we applied the same simulation of data generation and assimilation as that in our previous paper, which proposed the vbcEnKF scheme. By using the same procedure, we can directly compare the estimation performance with known exact solutions of E/I balance changes in synthetic observations.

In this simulation, we first generated 30 s time series of synthetic EEG data based on the NM model using the fourth-order Runge-Kutta method with the sampling interval $dt = 0.01$ using the following parameter conditions used previously [1]:

$$
\begin{aligned}
&A(t) = \begin{cases} 3.25\ (t \leq 10\text{s}) \\ 4.25\ (t > 10\text{s}) \end{cases} \\
&a(t) = 100 \\
&B(t) = \begin{cases} 22\ (t \leq 10\text{s}) \\ 19\ (t > 10\text{s}) \end{cases} \\
&b(t) = \begin{cases} 50\ (t \leq 10\text{s}) \\ 52\ (t > 10\text{s}) \end{cases} \\
&p(t) \sim \mathcal{N}(220,\ 22)
\end{aligned}
\tag{S1}
$$

Using such generated synthetic EEG data, we applied two proposed DA methods (based on either vbcEnKF or vbcETKF) 50 times with a different initial random seed for each ensemble size condition. The ensemble sizes of vbcEnKF and vbcETKF were selected from [40, 500] with 20 step size and [40, 150] with 10 step size, respectively. When applying the synthetic EEGs as observational data to each DA method, the EEGs were assimilated into the NM model sample-by-sample with the same sampling interval of the observation. Moreover, to reveal whether each DA method could correctly detect the parameter changes from observed data with noisy sampling, white noise was added to the synthetic EEGs using $\mathcal{N}(0,1.3)$. Afterwards, we evaluated the estimation error score of synthetic EEGs and the standard deviation of estimated noise covariance, which was obtained from 50 trials of the estimation for each ensemble size condition. The estimation error score was calculated using the mean absolute error (MAE) function described by the following equation:

$$
\text{MAE} = \frac{1}{N_t}\sum_{t=0}^{N_t}\left|y_{obs,t} - \hat{y}_t\right| \tag{S2}
$$

where $y_{obs,t}$ and $\hat{y}_t$ are the observed (exact) and predicted EEG at time sample $t$, respectively, and $N_t$ is the total number of samples. These evaluation scores are accessed for each DA method.

The results of this validation are shown in Figs. S1 and S2. Fig. S1 shows examples of state estimation results from the vbcEnKF-based (previous) method with ensemble size $N_{ens}$ = 200 and the vbcETKF-based (proposed) method with $N_{ens}$ = 70. Although the predicted time series of EEG and the mE/I ratio seemed to be qualitatively similar to the exact values between both DA methods (Fig. S1), $N_{ens}$ was different. Even though $N_{ens}$ in the proposed method vbcETKF is > 50% smaller than that in the previous method vbcEnKF, the proposed method can achieve similar estimation performance to that of the previous method. Indeed, as shown in Fig. S2a, b, the prediction error index (MAE) in the proposed method was around 0.2 points smaller than that of the previous method for all $N_{ens}$ conditions, and the proposed method exactly decreased the MAE score with small $N_{ens}$ relative to the previous method. Moreover, as shown in Fig. S2c, d, the previous method requires $N_{ens} \geq 200$ to estimate the exact observational noise covariance, whereas the proposed method can estimate observational noise covariance with $N_{ens} \geq 70$.

In summary, the proposed vbcETKF method could predict the model states and estimate parameters, and $N_{ens} \geq 70$ was required to guarantee an accurate prediction with a small prediction error of the observations. This indicates that the proposed method can achieve a > 50% decrease in $N_{ens}$ for the state and parameter estimation in the NM model with numerical simulations.

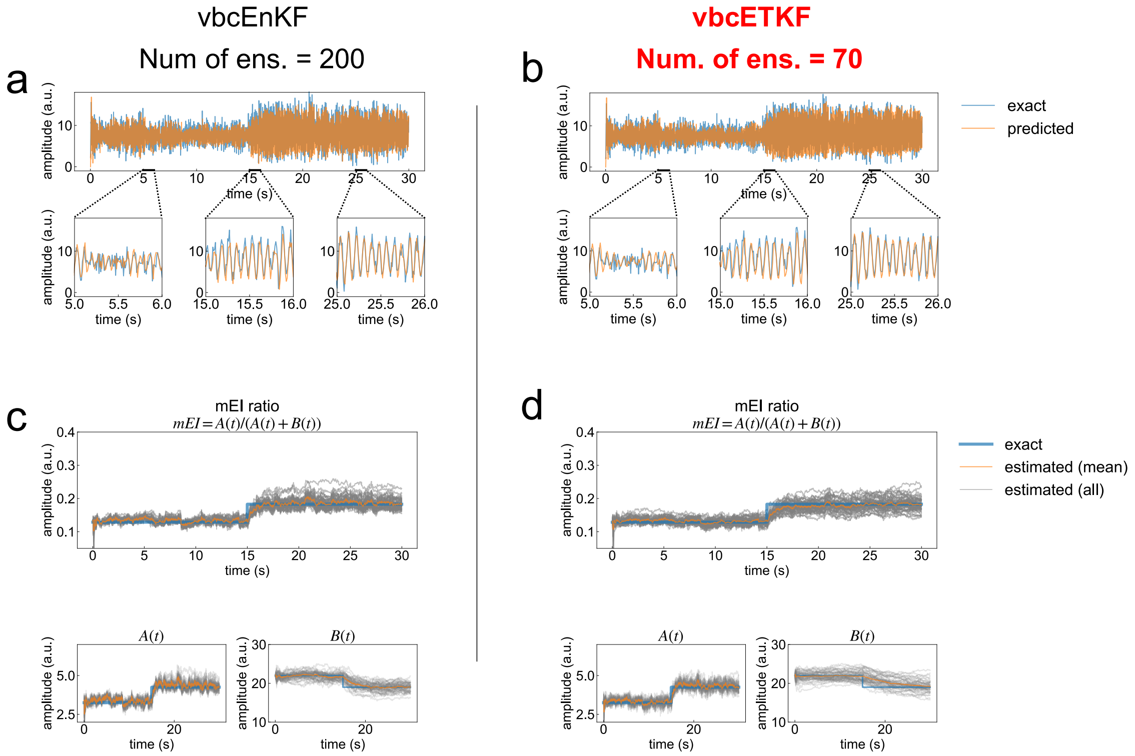


**Figure S1: Comparisons of time-series predictions for electroencephalographic signals (EEGs) and excitation/inhibition (E/I) balance changes.** Comparison between the original EEG data and typical predicted EEGs obtained from the first trial out of 50 trials with the vbcEnKF-based and vbcETKF-based methods. The upper panels in (**a**) and (**b**) show the entire time series of synthetic and predicted EEGs. The lower three panels in (**a**) and (**b**) are enlarged views of the prediction results for each time interval. Blue and orange lines indicate the exact and predicted EEG values, respectively. **c, d** Estimations of the model-based E/I (mE/I) ratio obtained from 50 trials with the vbcEnKF-based and vbcETKF-based methods. The upper panels in (**c**) and (**d**) show the synthetic and predicted mE/I ratios. The lower two panels in (**c**) and (**d**) show the synthetic and predicted excitatory and inhibitory gain parameters $A$ and $B$, respectively. Blue and orange lines indicate the time series of exact values and the mean of the estimations. Gray lines indicate the estimated value of each parameter obtained from 50 trials.

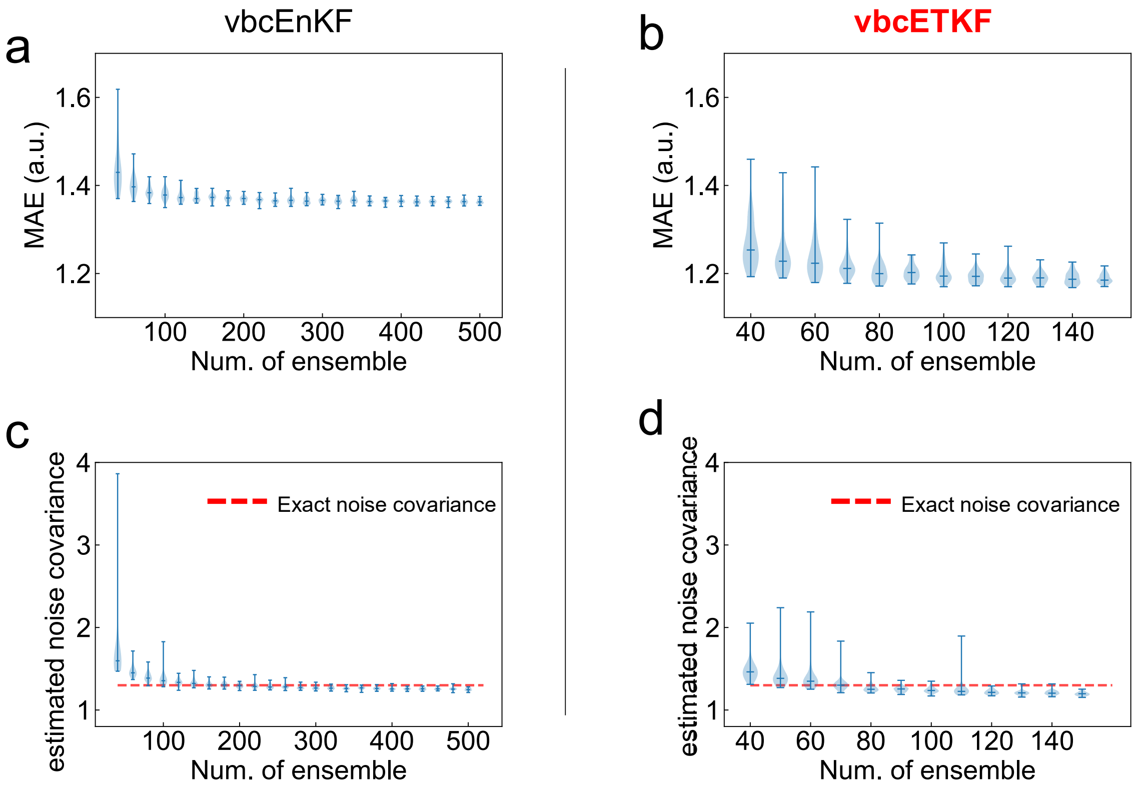


**Figure S2: Comparisons of the effects of ensemble size on estimation accuracy. a, b.** Violin plots of prediction error scores as a function of the number of ensembles obtained from the vbcEnKF-based (**a**) and vbcETKF-based (**b**) methods. The probability densities in these violin plots were estimated using a kernel density estimation method based on the samples obtained from 50 trials. **d, e**. Corresponding estimated observation noise covariance as a function of the number of ensembles obtained from the vbcEnKF-based (**c**) and vbcETKF-based (**d**) methods. The red dashed line indicates the exact noise covariance of synthetic electroencephalographic data. Error bars denote maximum and minimum values, and the middle lines denote median values. MAE: mean absolute error.

### *Supplementary References*


[1] Yokoyama, H. & Kitajo, K. A data assimilation method to track excitation-inhibition balance change using scalp eeg. *Communications Engineering* **2**, 92 (2023). URL https://www.nature.com/articles/ s44172-023-00143-7. https://doi.org/10.1038/s44172-023-00143-7.

[2] Burgers, G., Leeuwen, P. J. V. & Evensen, G. Analysis scheme in the ensemble kalman filter. *Monthly Weather Review* **126**, 1719–1724 (1998). https://doi.org/10.1175/1520-0493(1998)126⟨1719:ASITEK⟩2.0.CO;2.

[3] Whitaker, J. S. & Hamill, T. M. Ensemble data assimilation without perturbed observations. *Monthly Weather Review* **130**, 1913–1924 (2002). https://doi.org/10.1175/1520-0493(2002)130⟨1913:EDAWPO⟩2.0.CO;2.

[4] Bishop, C. H., Etherton, B. J. & Majumdar, S. J. Adaptive sampling with the ensemble transform kalman filter. part i: Theoretical aspects. *Monthly Weather Review* **129**, 420–436 (2001). URL http://journals.ametsoc.org/ doi/10.1175/1520-0493(2001)129⟨0420:ASWTET⟩2.0.CO;2. https://doi. org/10.1175/1520-0493(2001)129⟨0420:ASWTET⟩2.0.CO;2.
